# *In vitro* and *in vivo* validation of cwlM and pbpB essentiality for viability and resistance to imipenem in *Mycobacterium abscessus*

**DOI:** 10.1101/2024.03.08.584109

**Authors:** Jin Lee, Si-Yang Li, Dalin Rifat, Natalia Kurepina, Liang Chen, Barry N. Kreiswirth, Eric L. Nuermberger

## Abstract

*Mycobacterium abscessus* lung infection is notoriously difficult to treat due, in part, to the intrinsic resistance of this pathogen to most marketed antibiotics. β-Lactams, namely imipenem and cefoxitin, are first-line drugs in combination regimens used to treat this infection; and there is growing interest in dual-β-lactam-based regimens. Better understanding of the molecular basis of β-lactam activity through study of the genetic determinants of β-lactam susceptibility and tolerance would enable more rational drug combinations and guide discovery of novel drug targets to complement β-lactams. We recently used an inducible CRISPR interference (CRISPRi) system to silence *cwlM* and *pbpB* and confirm their essentiality for *in vitro* growth and resistance to sub-MIC concentrations of imipenem. Here, we extend those findings to show that silencing either gene alone is bactericidal and augments the bactericidal activity of imipenem *in vitro*. Furthermore, using CRISPRi in a mouse model of *M. abscessus* lung infection for the first time, we confirm the essentiality of each gene for *in vivo* survival. These results validate *cwlM* and *pbpB* as essential genes and promising drug targets in this pathogen, including for potentiation of carbapenem activity. The results further establish CRISPRi as a powerful method for validating drug targets and studying gene-gene and gene-drug interactions *in vitro* and *in vivo*.

## Introduction

Members of the *Mycobacterium abscessus* complex, including *M. abscessus* subsp. *abscessus*, are known for their intrinsic resistance to commonly used antibiotics and are increasingly recognized as evolving pathogens (1)(2). Pulmonary infections caused by *M. abscessus*, which typically occur in hosts with underlying structural lung disease and/or compromised immune responses, are particularly difficult to treat (3)(4)(5). They are essentially incurable without surgical excision and often result in a rapid deterioration of respiratory function (6). Recurrent cases account for more than 40-50% of the total cases (7)(8)(9) The poor clinical outcomes observed with recommended regimens indicate an urgent and pressing need for novel antibacterial drugs and treatment approaches. Despite the recent emergence of drug discovery and development programs directed at *M. abscessus*, the level of investment remains modest and there are many challenges, including the limited availability of qualified tools for target identification, *in vitro* and *ex vivo* validation, and animal models representative of human disease (10)(11).

Repurposing of antibiotics developed for other indications remains the major source of new drugs for *M. abscessus* pulmonary disease. β-Lactams are considered a cornerstone in currently recommended combination drug regimens; and there has been a growing interest in dual-β-lactam-based combination therapy. Several studies (12)(13)(14)(15) (16)(17)(18), including our own, have demonstrated synergistic effects when combining β-lactams with or without β-lactamase inhibitors against clinical *M. abscessus* isolates *in vitro*. This suggests that the use of β-lactam combinations holds promise in the treatment of *M. abscessus* infections. However, significant knowledge gaps exist that hinder the rational development of optimized β-lactam combination therapies for *M. abscessus* infections. First, the basis for selecting the optimal combination of β-lactam antibiotics remains unclear. Most studies have relied on random pairing of different β-lactam antibiotics and conducting *in vitro* susceptibility testing to identify effective combinations by chance. Second, most studies have focused on screening for growth inhibition *in vitro*, rather than bactericidal effects and *in vivo* efficacy. Better understanding of the molecular basis of β-lactam activity through study of the genetic determinants of β-lactam susceptibility and tolerance would enable more rational design of optimal combinations of two β-lactams, with or without a β-lactamase inhibitor, and of β-lactams with other drugs. Such study may also lead to identification of novel drug targets to fuel future drug discovery efforts.

We previously predicted the essentiality of 326 genes in the genome of the *M. abscessus* ATCC 19977 type strain using transposon insertion sequencing in a saturated transposon mutant library, identifying 3 predicted essential genes involved in peptidoglycan synthesis and remodeling (19). We subsequently used an inducible CRISPRi-based assay to conditionally silence the expression of two predicted essential peptidoglycan biosynthesis genes, *pbpB* and *cwIM*, in order to confirm their essentiality for *in vitro* growth (20). We also showed that silencing either gene sensitized *M. abscessus* to the growth-inhibitory effects of imipenem (IPM), a first-line β-lactam in recommended treatment regimens (20). These results demonstrated that CRISPRi provides an efficient experimental approach to study gene function and drug/target interactions in *M. abscessus*. In the current study, we performed time-kill assays to show that the loss of *pbpB* or *cwlM* does not just inhibit growth but is indeed lethal to *M. abscessus* and increases the bactericidal effects of IPM. We also demonstrated the utility of the CRISPRi platform for confirming gene essentiality *in vivo* by confirming that silencing of *pbpB* or *cwlM* is lethal in a mouse model of *M. abscessus* lung disease.

## Materials and methods

### Bacteria

*M. abscessus* subsp. *abscessus* strain ATCC 19977 was obtained from the American Type Culture Collection (ATCC) and used as the wild-type control. Strains were engineered for conditional silencing of *cwlM* and *pbpB* using the plasmid pLJR962, containing an ATc-inducible *dCas9*_*Sth1*_ and an ATc-inducible single guide RNA (sgRNA) along with a kanamycin selection marker, as the vector, as previously described (20). The empty vector control lacks the target sgRNA.

### Media

Bacterial cultures were initiated in standard growth media: Middlebrook 7H9 broth supplemented with 10% (v/v) Middlebrook OADC supplement, 0.1% (v/v) glycerol, and 0.05% (v/v) Tween 80. Drug activity assays in nutrient-rich media were performed using either Middlebrook 7H9 broth with 10% (v/v) OADC and 0.1% (v/v) glycerol but without Tween 80; or CAMHB without Tween 80. Polystyrene petri-dishes (100 mm × 15 mm) containing 20 mL 7H11 agar supplemented with 10% (v/v) OADC and 0.5% (v/v) glycerol were used to determine CFU counts. Difco BBL Mueller Hinton II broth (cation-adjusted) powder (i.e., CAMHB powder), Difco Middlebrook 7H9 broth powder, Difco Mycobacteria 7H11 agar powder, and BBL Middlebrook OADC enrichment were manufactured by Becton, Dickinson and Company. Glycerol and Tween 80 were purchased from Fisher Scientific. Selective 7H11 agar plates for enumerating CFU in lung homogenates contained with 50 μg/ml carbenicillin, 10 μg/ml polymyxin B, 20 μg/ml trimethoprim, and 50 μg/ml cycloheximide. Carbenicillin and cycloheximide was purchased from Research Products International, and polymyxin B and trimethoprim from Sigma-Aldrich.

### Drugs

Imipenem monohydrate (Biosynth International, Inch.) and kanamycin (Research Products International) were dissolved in water. Anhydrotetracycline (Sigma-Aldrich) was dissolved in dimethyl sulfoxide (DMSO). Dexamethasone (Sigma-Aldrich) was dissolved in normal saline before injection.

### Drug activity assays in nutrient-rich media

Frozen stocks of *M. abscessus* were cultured in standard growth media until the bacterial suspension reached OD_600_ ∼1 (approximately 10^7^-10^8^ CFU/mL), at which point the assays were initiated (“Day 0”) with approximately 5×10^5^ CFU/ml inoculated into each tube. Assays were conducted with the indicated media in a total volume of 2.5 mL in 14 mL, round-bottom, polystyrene tubes. MIC was determined using the broth macrodilution method, as previously described (21).

### Murine model of *M. abscessus* lung infection and inducible gene silencing *in vivo*

All animal procedures were approved by the Animal Care and Use Committee at Johns Hopkins University. 6-week-old female C3HeB/FeJ mice (Jackson Labs) were aerosol-infected with *M. abscessus* using the Inhalation Exposure System (Glas-Col) and a fresh log-phase broth culture (optical density at 600 nm of approximately 1.0) aiming for implantation of between 3.5 and 4.0 log_10_ CFU in the lungs of each mouse. Lungs from 3 control mice in each aerosol infection group were obtained one day after implantation to determine the number of bacteria in the lungs after each run. Dexamethasone was administered subcutaneously at 12.5 mg/Kg once daily for 4 days prior to infection and 3 days post-infection, then continued at a lower dose of 10 mg/Kg/d for the remainder of the experiment. Doxycycline (2000 parts per million) was provided in Purina rodent chow (Research Diets, Inc) (22), beginning on the day of infection to induce expression of the CRISPRi constructs. Thirteen days post-infection, mice from different treatment groups were sacrificed, and the lungs were homogenized and plated in serial 10-fold dilutions on selective 7H11 agar plates to determine the bacterial burden and response to treatment.

### Quantitative cultures, CFU counting, and analysis

In all assays, CFU counts were determined by culturing serial 10-fold dilutions of 0.5 ml bacterial suspensions in PBS on 7H11 or selective 7H11 agar containing 10% OADC. For *in vitro* drug activity assays, CFU counts were not determined for samples where the bacteria had overgrown, clumped, and fallen out of suspension. After all liquid was absorbed into the agar, the plates were sealed in plastic bags and incubated at 37°C for 7-10 days before the final reading of CFU counts. The dilution that yielded CFU counts between 10-120 and closest to 50 was used to determine CFU/mL. The CFU/mL value (*x*) was log transformed as log_10_ (*x* + 1) prior to analysis.

### Statistical analysis

Group mean CFU counts were compared using t test or one-way ANOVA with Dunnett’s post-test, respectively. GraphPad Prism 9 was used for all analyses.

## Results

In preparation for time-kill experiments to confirm the lethality of conditional silencing of *cwlM* and *pbpB*, the growth of each conditional knockdown (cKD) mutant was evaluated in CAMHB and 7H9 media supplemented with 50 μg/ml of kanamycin, with comparison to the ATCC 19977 parent strain and the empty vector control. No significant differences were noted between strains (data not shown).

The effects of conditional knockdown of *cwlM* and *pbpB* on viability were assessed in CAMHB media supplemented with kanamycin 50 μg/mL to maintain the CRISPRi construct. ATc concentrations ranging from 0.06 to 1 μg/mL had no effect on growth of the empty vector control, which formed pellets at each ATc concentration. In contrast, ATc restricted or prevented growth of each cKD mutant in a similar concentration-dependent fashion. Samples plated after three days of incubation with ATc 0.5-1 μg/mL demonstrated 0.35-0.50 log_10_ CFU reductions for each mutant compared to day 0. However, as ATc is known to be unstable in aqueous media at 37 deg C, the experiment was repeated with the cKD strains using a wider range of ATc concentrations (8 - 0.025 μg/mL) and daily supplementation of ATc by adding back 62.5% of the original concentration (23). Interestingly, with exposure to ATc concentrations ≤1 μg/mL for one day, conditional knockdown of *cwlM* was associated with greater CFU reductions than knockdown of *pbpB* (**Fig. 1, left panel**). However, after three days of ATc exposure, conditional knockdown of *pbpB* was more lethal because CFU counts of the cKD-*pbpB* mutant had continued to decline, while CFU counts of the cKD-*cwlM* actually increased between day 1 and day 3 in tubes containing ATc concentrations ≤1μg/mL (**Fig. 1, right panel**).

**Fig. 1.**
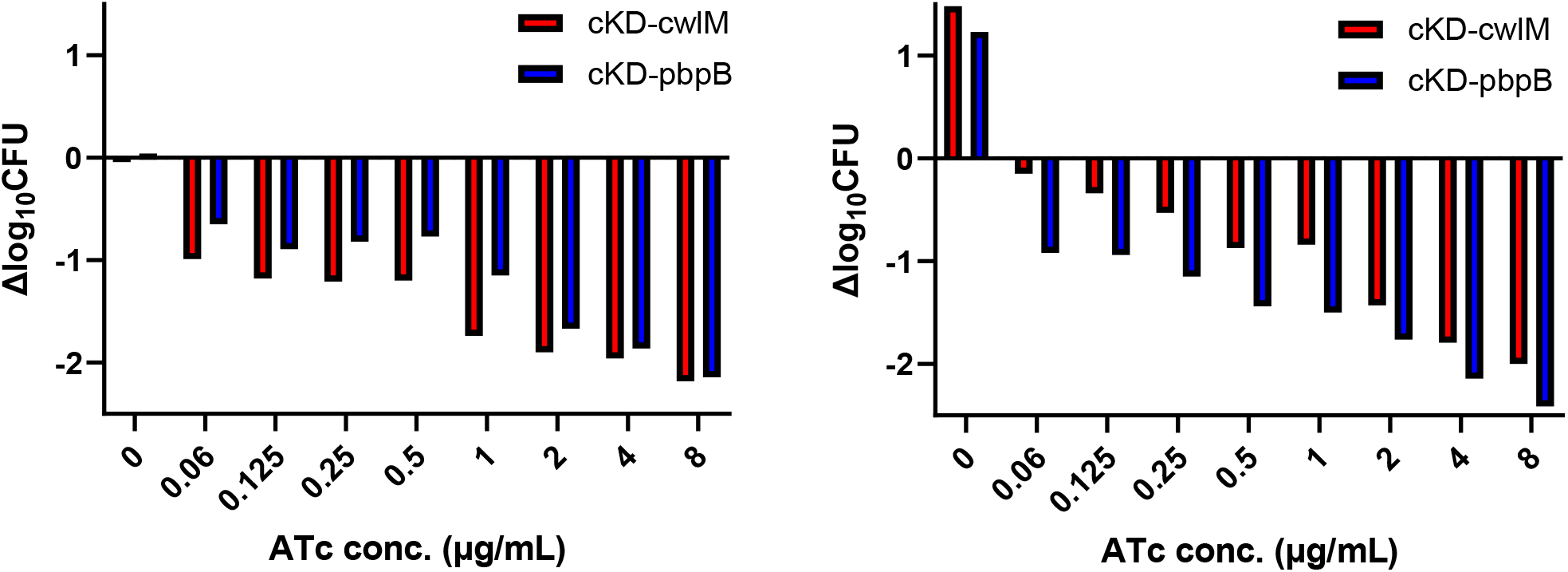
Time-kill study with exposure of cKD mutants to increasing ATc concentrations. ATc was supplemented daily, at 62.5% of original concentration. Results after 1 day (left) and 3 days (right) of incubation are presented as the change in log_10_ CFU/mL compared to the CFU count at Time 0.

We previously reported that silencing of *cwlM* and *pbpB* expression sensitized *M. abscessus* to growth inhibition by IPM (20). Here, we assessed the effects of conditional knockdown of *cwlM* and *pbpB* on the bactericidal activity of IPM in CAMHB. In CAMHB without ATc, bactericidal effects were observed only at IPM concentrations ≥4 μg/mL (**Fig. 2, top left panel**). Neither cKD mutant was more susceptible to IPM than the empty vector control in the absence of ATc. When the experiment was repeated with plating of the drug-free control, with addition of kanamycin in the media and in the agar plates at 50 μg/mL, the strains again showed no difference in susceptibility to IPM in the absence of ATc (**Fig. 2, top right panel**). The IPM MIC was 4 μg/mL against all strains.

**Fig. 2.**
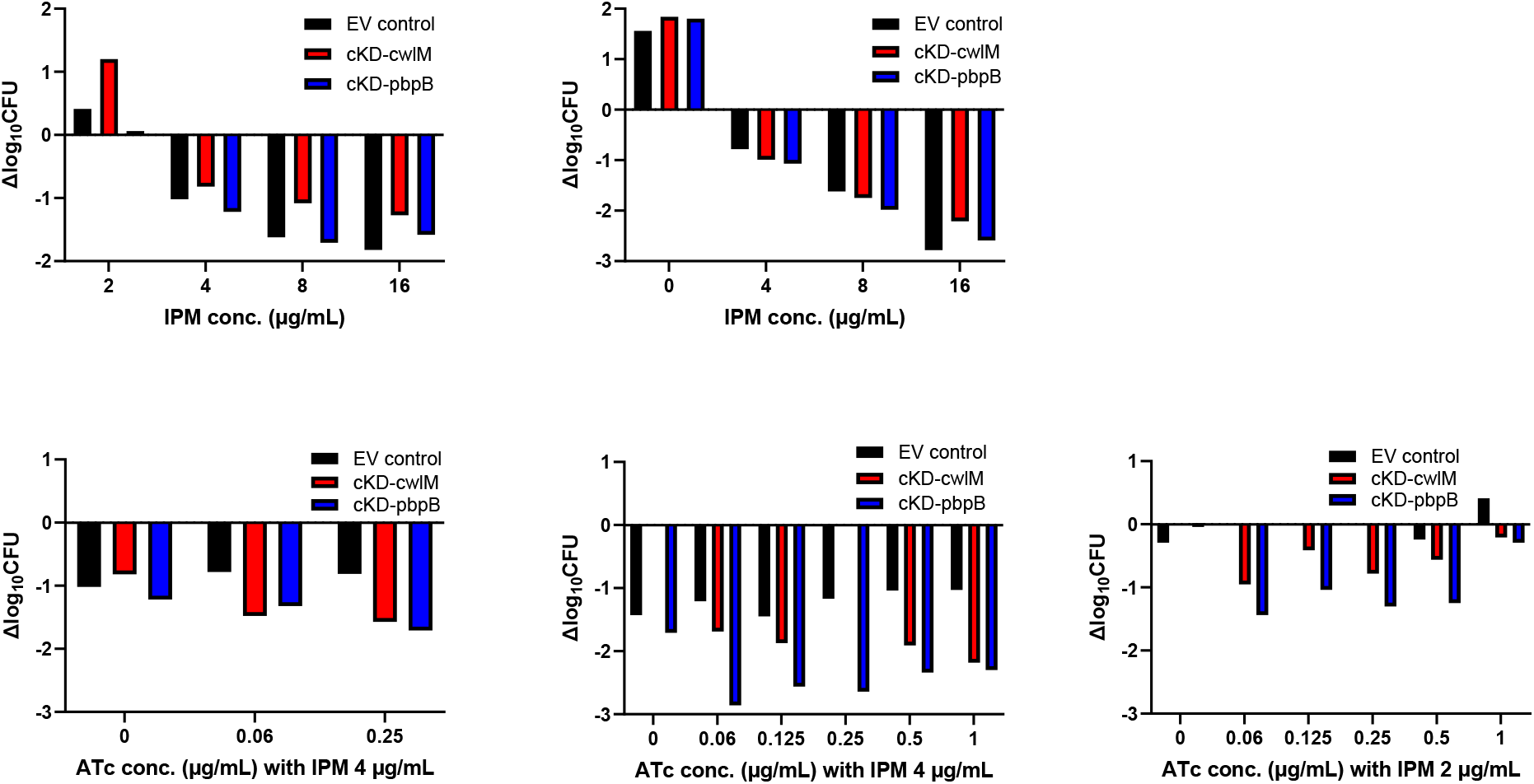
Time-kill study with exposure of the empty vector (EV) control and cKD mutants to increasing IPM concentrations in the absence (top row) and presence (bottom row) of ATc induction. The concentration response for IPM alone is shown in the absence (top left) and presence (top right) of selective kanamycin (50 μg/mL). The effect of increasing concentrations of ATc on IPM activity is shown without (bottom left) and with (bottom middle and bottom right) daily (62.5%) ATc supplementation. Results are after 3 days of incubation and presented as the change in log_10_ CFU/mL compared to the CFU count at Time 0.

With addition of ATc at 0.06 or 0.25 μg/mL, both mutants were modestly more susceptible to bactericidal effects of IPM at 4 μg/mL, while the empty vector control was not (**Fig. 2, bottom left panel**). The effects of silencing *cwlM* and *pbpB* with ATc in the presence of IPM at 4 μg/mL were roughly similar to the effects of increasing the IPM concentration from 4 μg/mL to 8-16 μg/mL in the absence of ATc. When susceptibility to IPM at 4 μg/mL was re-assessed with supplementation of ATc by adding back 62.5% of the original ATc concentration daily, the effects of silencing *cwlM* and especially *pbpB* on IPM susceptibility were even greater (**Fig. 2, bottom middle panel**). When IPM was used at 2 μg/mL, its effect was bacteriostatic, while, silencing *cwlM* and *pbpB* with daily supplementation of ATc was again bactericidal (**Fig. 2, bottom right panel**).

### *In vivo* validation of essentiality

Two experiments were performed in an aerosol infection model of *M. abscessus* lung disease in dexamethasone-treated C3HeB/FeJ mice to confirm the essentiality of *cwlM* and *pbpB in vivo*. In each experiment, mice were infected with either cKD mutant (*cwlM* or *pbpB*) or a control strain. For half of the mice in each arm, conditional silencing was induced using doxycycline provided in the mouse chow, beginning on the day of infection. The remaining mice received chow without doxycycline. In the first experiment, control mice were infected with the rough morphotype of the ATCC 19977 strain previously used to develop the mouse model. In the second experiment, control mice were infected with the empty vector control. In the first experiment, the ATCC 19977 rough morphotype control grew robustly, with median lung CFU counts increasing by more than 5 log_10_ in the presence or absence of doxycycline treatment (**Fig. 3, top left panel**). Likewise, the cKD-*pbpB* mutant CFU counts increased by approximately 4 log_10_ in mice that did not receive doxycycline (**Fig. 3, top middle panel**). However, with doxycycline treatment, the median CFU count on kanamycin-containing plates decreased by more than 2 log_10_ compared to day 0. The cKD-*cwlM* mutant did not grow as robustly in the absence of doxycycline, with median CFU counts increasing by only 2 log_10_ (**Fig. 3, top right panel**). However, with doxycycline treatment, median CFU counts on kanamycin-containing plates were again more than 2 log_10_ lower than day 0 counts. In the second experiment, the empty vector control strain increased by approximately 3 log_10_ CFU in mice with or without doxycycline treatment (**Fig. 3, bottom left panel**), as did the cKD mutants in the absence of doxycycline treatment. With doxycycline treatment, the median CFU counts of the *pbpB* and *cwlM* cKD mutants were approximately 3 log_10_ and 4 log_10_ lower than day 0 counts in the presence of doxycycline treatment. Taken together, the results of these experiments demonstrate the feasibility of conditional gene knockdown *in vivo* using CRISPRi constructs and confirm the *in vivo* essentiality of *cwlM* and *pbpB*, as demonstrated by the marked loss of viability with conditional silencing of either gene.

**Fig. 3.**
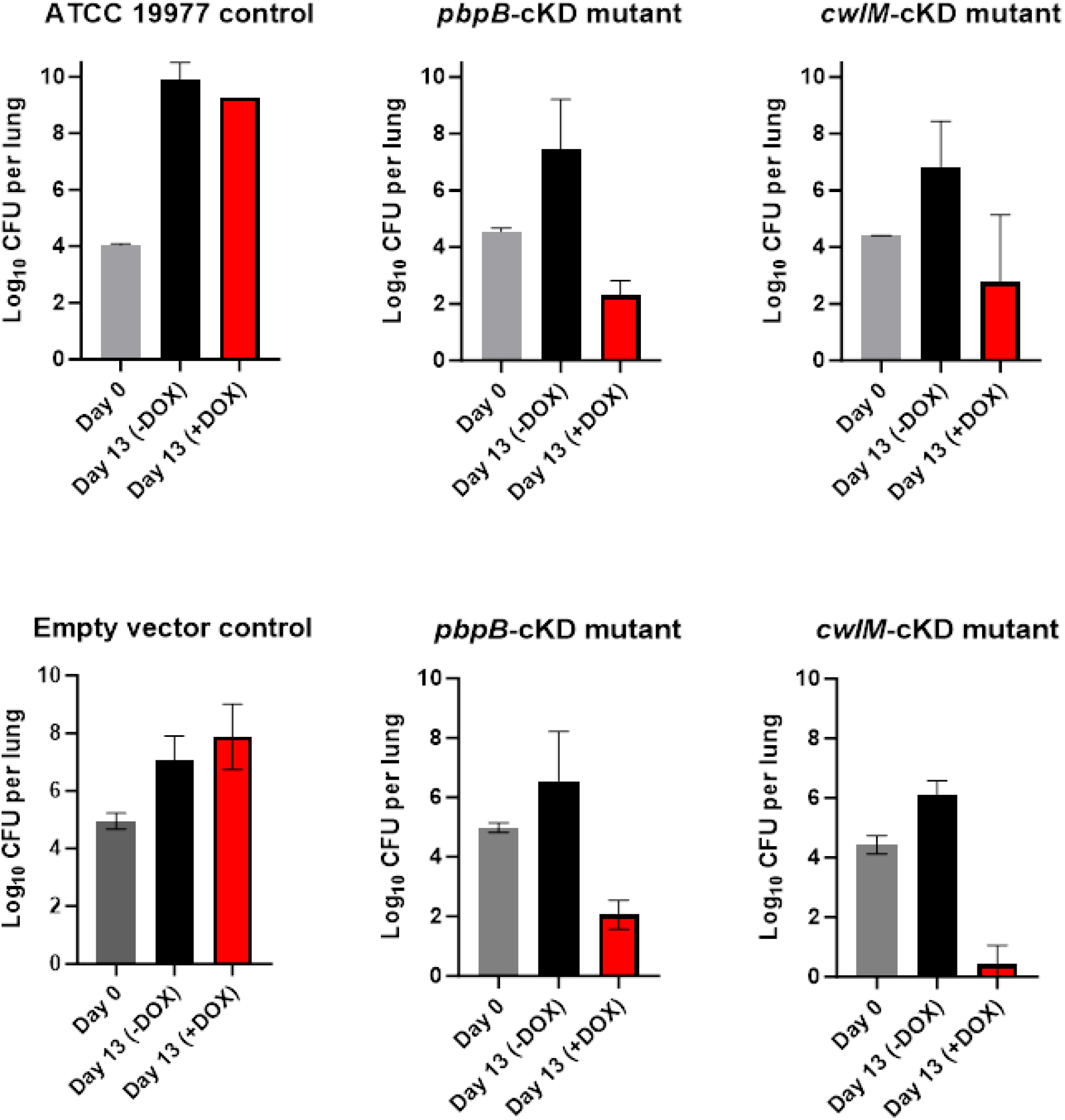
Loss of viability with doxycycline-induced silencing of *pbpB* and *cwlM* in a C3HeB/FeJ mouse model of *M. abscessus* lung infection. Results of two independent experiments are shown.

## Discussion

The results presented here confirm the essentiality of two peptidoglycan biosynthesis genes, *pbpB* and *cwIM*, for the viability of *M. abscessus*, as previously predicted by transposon insertion sequencing of our saturated transposon mutant library (19). The results extend our previous findings (20) to show that silencing the expression of either gene not only prevented growth and sensitized *M. abscessus* to the growth-inhibitory effects of IPM, but that such silencing was bactericidal and increased the bactericidal effects of IPM. As such, the results also support the hypothesis that these essential bacterial proteins may serve as promising therapeutic targets in *M. abscessus*. Our results add to initial studies by ourselves and others (11)(20)(29) to demonstrate that inducible CRISPRi platforms provide an efficient experimental approach to study conditional gene essentiality in *M. abscessus* and, importantly, extend the approach to the study of gene-drug interactions. More specific to drug discovery and regimen development applications, the platform can be useful for understanding molecular mechanisms of intrinsic antibiotic resistance and drug synergy, validating new drug targets including those whose inhibition is synergistically lethal with existing drugs, and screening for new chemical inhibitors of a specific target protein or pathway.

Despite increasing interest in dual-β-lactam combinations for treatment of *M. abscessus* infection, it remains unknown which minimal target sets need to be inhibited to overcome target redundancy and result in additive or synergistic β-lactam combinations (12)(24)(25). In contrast to the current testing approaches to mix and match different β-lactams, synergistic gene-gene and gene-β-lactam interactions can be used to develop and confirm hypotheses related to minimal target sets (and specific β-lactams) needed to overcome target redundancy. This strategy supports the rational selection of drug combinations or novel targets for drug discovery based on evidence that inhibition would potentiate existing drugs. Whether silencing of *pbpB* potentiates IPM activity because PbpB is a binding target of IPM or because PbpB is able to compensate for loss of another IPM binding target (or both) will require further study. PbpB localizes to the septum and is largely responsible for septal PG synthesis (26)(27)(28). Recent work has demonstrated synergistic lethality from simultaneous knockdown of *pbpB* and the gene encoding another essential PBP in *M. abscessus*, PBP-lipo, and that overexpression of *pbpB* rescues the growth defect and morphological changes caused by knockdown of PBP-lipo, indicating that these proteins have overlapping functions at the septum (29). Meropenem and faropenem were previously shown to covalently acylate the *M. tuberculosis* homolog of PbpB (30) but, to our knowledge, it is not known whether PBP-lipo is a binding target of carbapenems and penems in either mycobacterial species.

Initially annotated as an amidase, CwlM is likely a key regulator of peptidoglycan synthesis, as shown in *M. tuberculosis* (31), where it regulates biosynthesis of peptidoglycan precursors and their transport across the cytoplasmic membrane. It is unlikely to be a binding target of IPM. More likely is that silencing of *cwlM* depletes *M. abscessus* of critical precursors for peptidoglycan synthesis to reduce substrate availability for the PBPs and LDTs that are targeted by IPM and potentially any compensatory transpeptidase enzymes that are not targeted effectively by IPM. Therefore, our data further reinforce CwlM as a unique target in its own right, for which an inhibitor would also be expected to potentiate β-lactam activity against *M. abscessus*.

No consensus exists on the most appropriate nonclinical models for evaluating drug efficacy against *M. abscessus* lung infection. Immunocompetent mouse strains are not susceptible to persistent *M. abscessus* infection. Limited experience indicates that immunocompromised athymic nude or SCID mice maintain sufficient bacterial burdens to assess treatment efficacy over short durations, but they are expensive and do not develop necrotic lung lesions characteristic of advanced *M. abscessus* lung disease (32)(33)(34). We developed a progressive *M. abscessus* lung infection model in dexamethasone-treated C3HeB/FeJ mice that produces high lung CFU burdens and can develop more representative necrotizing granulomatous lung lesions (35)(36). Here, we use this model to demonstrate the utility of the inducible CRISPRi platform for confirming gene essentiality *in vivo* by demonstrating that conditional silencing of *pbpB* or *cwlM* is lethal during *in vivo* infection. To our knowledge, this is the first published use of inducible CRISPRi in a mouse model of *M. abscessus* lung disease. Although this approach may be most useful for approximating the effects of inhibiting essential drug targets, for which knockout mutants cannot be generated, it is also useful for testing hypotheses related to genes that are non-essential for growth but are conditionally essential in the presence of one or more antibiotics and/or under specific environmental conditions. The murine experiments presented here were designed to show proof-of-concept for *in vivo* gene essentiality irrespective of the development of necrotic lung lesions. Therefore, induction of CRISPRi with doxycycline was implemented shortly after infection and dexamethasone was continued for the duration of the experiments. We acknowledge that these conditions are unlikely to replicate the conditions of an established, persistent lung infection with *M. abscessus*.

